# Learning in anticipation of reward and punishment: Perspectives across the human lifespan

**DOI:** 10.1101/738211

**Authors:** Matthew J. Betts, Anni Richter, Lieke de Boer, Jana Tegelbeckers, Valentina Perosa, Rumana Chowdhury, Raymond J. Dolan, Constanze Seidenbecher, Björn H. Schott, Emrah Düzel, Marc Guitart-Masip, Kerstin Krauel

## Abstract

Pavlovian biases influence the interaction between action and valence by coupling reward seeking to action invigoration and punishment avoidance to action suppression. In this study we used an orthogonalised go/no-go task to investigate learning in 247 individuals across the human lifespan (7-80 years) to demonstrate that all participants, independently of age, demonstrated an influence of Pavlovian control. Computational modeling revealed peak performance in young adults was attributable to greater sensitivity to both rewards and punishment. However in children and adolescents an increased bias towards action but not reward sensitivity was observed. In contrast, reduced learning in midlife and older adults was accompanied with decreased reward sensitivity and especially punishment sensitivity. These findings reveal distinct learning capabilities across the human lifespan that cannot be probed using conventional *go/reward no-go/*punishment style paradigms that have important implications in life-long education.

## Introduction

Adaptive behavioral choices maximize reward and minimize punishment. In order to achieve this optimization, humans are equipped with two broad classes of mechanisms. Firstly, a Pavlovian controller directly ties affectively important outcomes together with learned predictions of these outcomes, to valence-dependent stereotyped behavioral responses via a conceived hard-wired mechanism. Secondly, a more flexible, instrumental controller learns choices on the basis of contingent consequences (Dayan and Balleine, 2002). Generally these controllers favor the same choices rendering learning fast and efficient. However in some circumstances, Pavlovian influences may impair instrumental learning by prescribing the opposite of the instrumental controller leading to sub-optimal behavioral choices (Dayan *et al.*, 2006; Guitart-Masip *et al.*, 2010).

A number of studies have shown that the success of instrumental learning critically depends on whether reward or avoidance of punishment is paired with action or inhibition. Using a go/no-go task that independently dissociates, i.e. orthogonalizes, action and valence, young adults demonstrate striking asymmetry in instrumental learning by being better at learning to emit a behavioral response in anticipation of reward and better at withholding a response in anticipation of punishment (Guitart-Masip *et al.*, 2012*b*; Cavanagh *et al.*, 2013, Chowdhury *et al.*, 2013*b*; Guitart-Masip *et al.*, 2014; Richter *et al.*, 2014). Computational modeling approaches in young adults have shown this asymmetry in instrumental action learning is due to a Pavlovian coupling between action and valence expectation where the strength of this bias relates to failure in learning the conflicting conditions: withholding a response in anticipation of reward and emitting a response in anticipation of punishment (Guitart-Masip *et al.*, 2012*b*; Cavanagh *et al.*, 2013; Guitart-Masip *et al.*, 2013). However a recent study has shown that this coupling may also be due to instrumental mechanisms (Swart *et al.*, 2017). Learning success in this go/no-go task requires flexibility, inhibition and the ability to use feedback and to detect reward contingencies. All of these abilities may critically rely on prefrontal cortex (PFC) dependent executive functions, which develop during adolescence but also decline in older age (Kray *et al.*, 2004; Zelazo *et al.*, 2004). Whilst most of the previous studies using the aforementioned task have investigated young adults, little attention has focused on how the processes underlying instrumental learning and potential conflict with Pavlovian control may change during differential periods across the lifespan— namely during childhood and adolescence, young, midlife and older age.

It has been widely stated that adolescents are highly sensitive to reward, which may contribute to increased risky behavior during this developmental period (Casey *et al.*, 2008). However it has also been suggested that this reward sensitivity may be adaptive by promoting learning and exploration — critical for transition into adulthood (Spear, 2000; Casey, 2015). A recent study demonstrated that adolescents learn to preferentially seek rewards rather than to avoid punishments, whereas young adults learn both behaviors equally well (Davidow *et al.*, 2016). However, previous studies have not dissociated reward sensitivity from action learning, and it remains thus unclear if this interpretation may be confounded by action requirements or to what extent changes in reward sensitivity may influence the strength of coupling between action and valence. This is particularly relevant in light of differential functional and anatomical development of limbic regions, such as the striatum and cognitive control regions during adolescence (Blakemore and Robbins, 2012; Shulman *et al.*, 2016). Such, asymmetrical development may translate into differential Pavlovian and instrumental strategies used by children and adolescents compared to those employed in adulthood.

The human brain also undergoes substantial change during normal aging, which has been associated with numerous cognitive changes (Bäckman *et al.*, 2006; Lindenberger, 2014). However it is not known how age-related differences in Pavlovian and instrumental control may impact behavioral inflexibility in older adults or whether such age-related changes are already evident in midlife adulthood. Previous work has shown that administration of the dopamine precursor L-DOPA enhances the Pavlovian influence of potential reward (Rutledge *et al.*, 2015) and may restore reward prediction errors in old age (Chowdhury *et al.*, 2013*a*). Coupled with the known age-related decline in the integrity of the dopaminergic system (Karrer *et al.*, 2017), a loss of functional dopamine may lead to a decrease in Pavlovian control in older age. Alternatively, previous studies have shown that the PFC is involved in overcoming the Pavlovian bias in young adults (Guitart-Masip *et al.*, 2012*b*; Cavanagh *et al.*, 2013). Thus, decreased functionality of the PFC as a result of normal aging could also lead to increased Pavlovian biases in older adults.

The objective of this study was to explore how acquisition of optimal/adaptive behavioral choices is differentially altered across the lifespan using an established go/no-go task that orthogonalizes action and valence. In addition we used computational modeling to investigate whether Pavlovian congruency is already evident in healthy children and adolescents in support of a hard-wired bias and compared behavior to young, midlife and older adults. Finally we assessed the extent to which action and valence are coupled across the lifespan and whether flexible learning to orthogonalize these two axes of behavioral control may vary with age.

## Results

247 healthy individuals from four age groups, namely children and adolescents (age: 7 to 16, n = 68), young adults (age: 18 to 30, n = 77), midlife adults (age: 31 to 60, n = 58), and older adults (age 61 to 80, n = 44) performed a previously established valenced go/no-go probabilistic learning task (Guitart-Masip *et al.*, 2012*b*). Participants had to learn through trial and error which of four fractal cues, preceding an easy visual target detection task, indicated the need (1) to respond to obtain a monetary reward (*go to win*), (2) to respond to avoid a monetary loss (*go to avoid losing*), (3) to withhold a response to obtain a monetary reward (*no-go to win*) and (4) to withhold a response to avoid a monetary loss (*no-go to avoid losing*) (see Figure 1). The outcome was probabilistic whereby correct feedback was given in 80% of win/loss conditions. To characterize the interaction of action and valence across the lifespan we investigated acquisition of and overall accuracy in each of the four conditions and analyzed differential influences of Pavlovian and instrumental learning using computational modeling.

**Figure 1:**
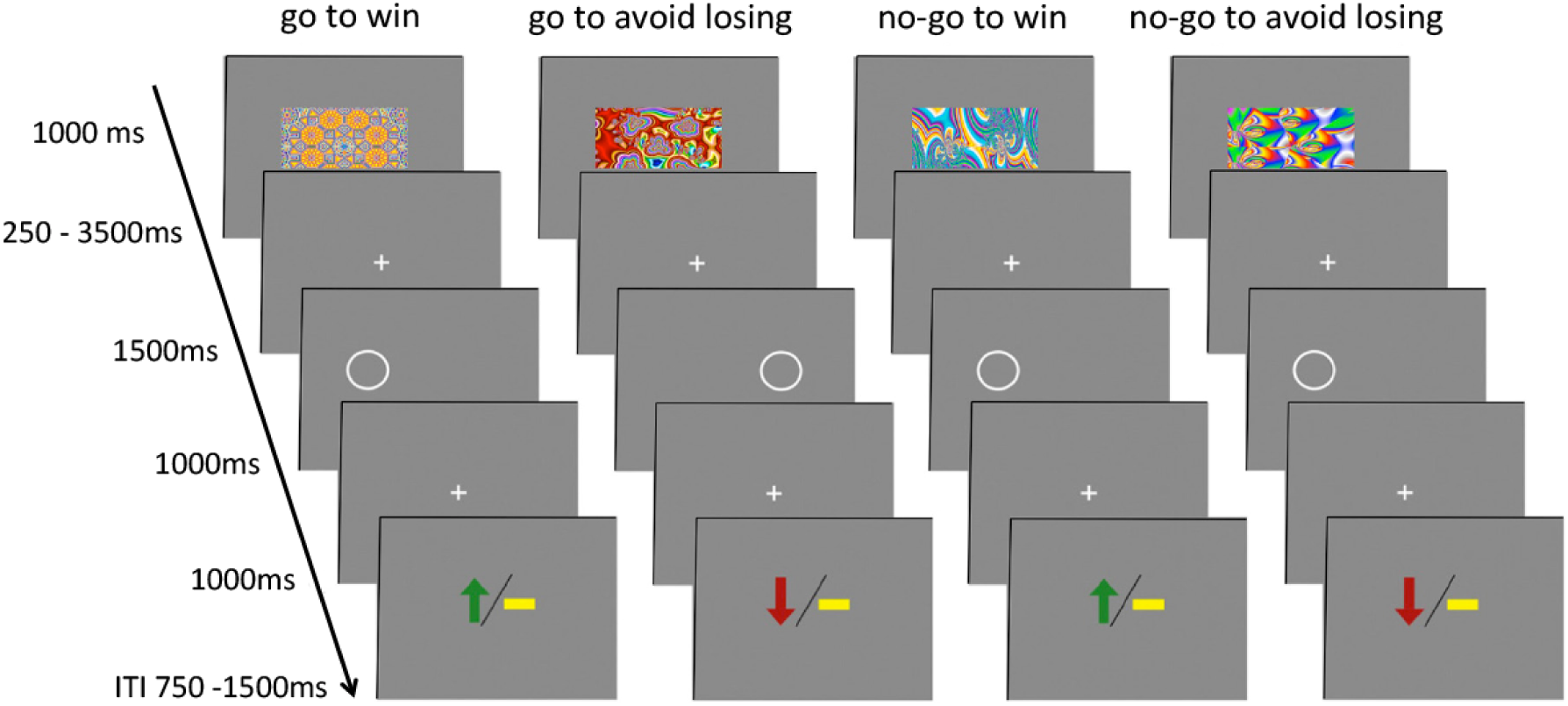
Probabilistic monetary go/no-go task. Participants had to learn over the course of 60 trials per condition, which fractal image was associated with responding or withholding a response to achieve a successful outcome (win or avoid losing). Responses indicated whether the circle presented after the fractal was located on the right or left side. Correct feedback was only provided in 80% of the trials. Abbreviations: ITI – inter-trial interval.

### Pavlovian bias across the lifespan

Performance as defined by percentage of correct (optimal) choices was assessed using a 4-way analysis of covariance (ANCOVA) with action (go/no-go), valence (win/lose) and time (1^st^/2^nd^ half) as within-subject factors, age group (children and adolescents/young adults/midlife/older adults) as a between-subject factor and gender as a covariate. As in previous studies (Guitart-Masip *et al.*, 2012*b*, 2014; Richter *et al.*, 2014) we observed main effects of time (F_1,243_ = 17.18, p < 0.0001) and action (F_1,243_ = 38.67, p < 0.0001) as well as an action x time interaction (F_1,243_ = 8.27, p =0.004), and an action x valence interaction (F_1,243_ = 14.21, p < 0.0005). Subjects demonstrated an increase in performance from the first to the second half of the experiment (*t*_*246*_ = -14.90, *p* < 0.0001) and performed better in conditions requiring a *go* choice than in trials requiring a *no-go* choice (*t*_246_ = 15.22, *p* < 0.0001). Participants demonstrated an initial bias toward *go* responses (1^st^(*go-nogo*)>2^nd^(*go-nogo*): *t*_246_ = 7.23, *p* < 0.0001), and showed greater accuracy for go choices when the outcome was a reward (*go to win > go to avoid losing: t*_246_ = 10.45, *p* < 0.0001) and for no-go choices when the outcome was avoidance of losses (*no-go to avoid losing > no-go to win: t*_246_ = 7.82, *p* < 0.0001).

Of particular interest in this study, were age-related differences in performance across groups (see Figure 2 and SI Table 1 for statistics). We observed a main effect of group (*p* < 0.0001), and interactions of group with action (*p* < 0.0001), valence (*p* = 0.033) and time (*p* = 0.001). Moreover a threefold interaction of action x time x group (*p* = 0.040) was found demonstrating a differential ability to learn action responses across the lifespan. However, no action x valence x group interaction was observed (F_3,243_ = 2.08, *p* = 0.10) suggesting human learning and decision making is influenced by Pavlovian control irrespective of age (Figure 2). No significant interactions with gender were observed (all p ≥ 0.2).

**Table 1:** Integrated Bayesian Information Criterion (iBIC) for all tested models

| Model no. | Model parameters | No. of parameters | Likelihood | Pseudo-R<sup>2</sup> | iBIC |
| --- | --- | --- | --- | --- | --- |
| 1 | $\epsilon, \rho$ | 2 | -28166 | 0.32 | 56,376 |
| 2 | $\epsilon, \rho, \xi$ | 3 | -27977 | 0.32 | 56,019 |
| 3 | $\epsilon, \rho, \xi, b$ | 4 | -25653 | 0.38 | 51,394 |
| 4 | $\epsilon, \rho_{\text{win}}, \rho_{\text{lose}}, \xi, b$ | 5 | -25064 | 0.39 | 50,239 |
| 5 | $\epsilon, \rho_{\text{win}}, \rho_{\text{lose}}, \xi, b, \pi_{\text{fluct}}$ | 6 | -24680 | 0.40 | 49,493 |
| **6** | **$\epsilon, \rho_{\text{win}}, \rho_{\text{lose}}, \xi, b, \pi_{\text{constant}}$** | **6** | **-24519** | **0.41** | **49,170** |

The winning model statistics are highlighted in bold font: $\epsilon$ , learning rate; $\rho_{\text{win}}$ , weighting of reward on win trials; $\rho_{\text{lose}}$ , weighting of punishments on lose trials; $b$ , go bias; $\pi$ , Pavlovian bias; $\xi$ , irreducible noise. iBIC, integrated Bayesian information criterion. Smaller values indicate a better model fit.

**Figure 2:**
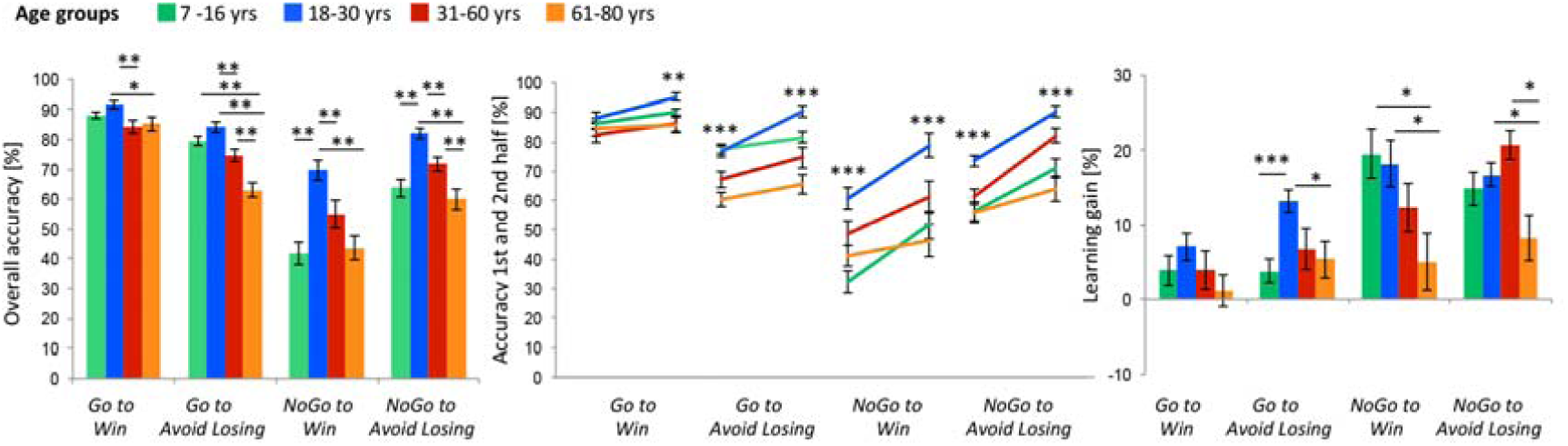
Overall behavioral performance across age groups. On the left, mean performance accuracy (%) ± standard error (SEM) across conditions in children and adolescents (7 to 16 years, n=69), young adults (18 to 30 years, n=77), midlife adults (31 to 60 years, n=58) and older adults (61 to 80 years, n=44) for all trials. In the middle panel, line graphs show mean performance for responses in the first (30 trials) and second half of trials (last 30 trials) across groups and on the right, bar plots demonstrate learning gain between first and second 30 trials across groups. ***(p<0.001), **(p<0.001), *(p<0.008; Holm-Bonferroni-corrected for 6 tests), indicate significant differences compared to young adults (independent samples t-tests).

To compare differences in performance across age groups, we performed independent samples t-tests and applied Holm-Bonferroni correction for six tests. We focused on overall accuracy, initial performance (first half) and ability to learn across trials (learning gain (2^nd^ - 1^st^ half) as shown in SI Table 1.

Young adults showed superior overall performance across all conditions compared to any other age group (all p<0.0001) and higher learning gain compared to older adults (p<0.0001). In *go trials*, children and adolescents demonstrated comparable performance to young adults whilst midlife and older adults demonstrated a significant age-related decline in *go* accuracy (all p<0.001). However, children and adolescents demonstrated significantly reduced learning gain for *go* trials compared to young adults (p=0.001). In *no-go* trials, young adults demonstrated superior overall performance compared to all other age groups irrespective of valence (all p<0.001). Taken together, children and adolescents demonstrated high performance in go trials comparable to young adults but not in no-go trials, indicating a predominant preference for action over withdrawal responses. In comparison, midlife and older adults demonstrated a significant age-related decline in both go and no-go trials compared to younger adults.

Interestingly assessment of learning gain revealed that children and adolescents were able to overcome this initial action bias through learning, demonstrating comparable learning gain to young adults for no-go trials (Figure 2). Midlife adults also showed poor initial *no-go* accuracy compared to younger adults (all p<0.001), but could increase performance in the second half of the experiment, significantly more so than older adults (p=0.001). In comparison, older adults demonstrated poor initial *no-go* accuracy (all p≤0.007) compared to young and midlife adults and considerable inflexibility in learning no-go responses across trials compared to all other groups (all p<0.004). Lastly older adults demonstrated the worst performance in punishment trials compared to all other age groups (all p≤0.001).

For completeness, reaction times for *go* responses are reported in the supplementary information (SI Figure 1). Their interpretation warrants caution as participants were explicitly instructed to respond accurately, while speed was not emphasized. Generally young and older adults demonstrated the fastest and slowest responses respectively (all p<0.0001) while children/adolescents and midlife adults did not differ (p=1.0).

### Parameterizing learning and biases using computational modeling

To identify instrumental and Pavlovian components of the observed asymmetry during learning, six nested reinforcement learning (RL) models were fitted to the behavioral data (see SI Materials and Methods) using the expectation maximization approach as previously described (Huys *et al.*, 2011, Guitart-Masip *et al.*, 2012*b*). All six computational models were fit to the data using a single distribution for all participants. This fitting procedure was, therefore, blind to the existence of different groups with putatively different parameter values. Our computational modeling approach demonstrated that the marked asymmetry in learning (i.e. superior performance in *go to win and no-go to avoid losing* compared to *go to avoid losing and no-go to win*) could be attributed to an interaction between instrumental and Pavlovian control mechanisms (Guitart-Masip *et al.*, 2012*b*). The best account of the data was provided by the model including a *static action bias, Pavlovian bias, reward and punishment sensitivity, learning rate* and an *irreducible noise* parameter consistent with previous studies using this task (see Table 1) (Guitart-Masip *et al.*, 2012*b*; Cavanagh *et al.*, 2013; Guitart-Masip *et al.*, 2014).

We found associative learning in children and adolescents was inherently driven by an action bias that was significantly greater than that observed in all adult age groups (Kruskal-Wallis with Wilcoxon rank-sum *post-hoc* tests; χ^2^ (3) = 15.42, all p ≤ 0.01, children and adolescents > young adults, midlife adults, older adults). Effect sizes determined using Cohen’s *d* between children/adolescents and all adult age groups were d ≥ 0.5. Parameters indicative of instrumental learning such as *sensitivity to reward* (χ^2^ (3) = 41.37; p < .0001), *sensitivity to punishment* (χ^2^ (3) = 39.33; p < 0.0001) and *learning rate* (χ^2^ (3) = 18.37, p < 0.001) followed an inverted U-shaped distribution, peaking during young adulthood (Figure 3). Children and adolescents demonstrated comparable *sensitivity to reward, punishment* and *learning rate* compared midlife adults (all p≥ 0.2) but increased *sensitivity to reward and punishment* (Wilcoxon rank-sum *post-hoc* tests; both Z ≤ -3.35, p ≤ 0.001; effect size d ≥ 0.5) but not *learning rate* (p=0.67) compared to older adults.

**Figure 3:**
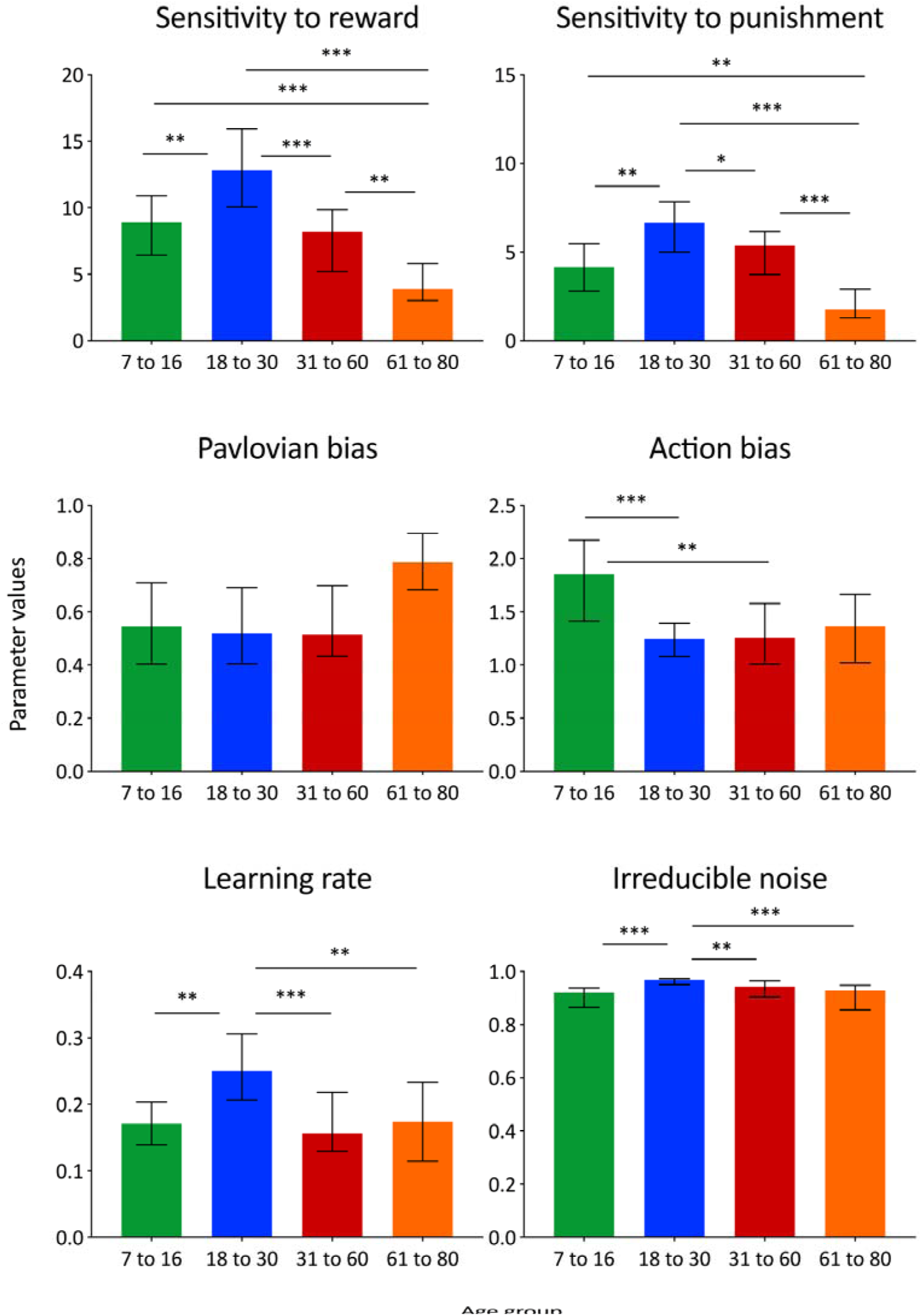
Modeled behavioural performance across the lifespan. Modeling parameters (median (5-95^th^ percentile)) derived from the winning model plotted across age groups. ***(p<0.001), **(p<0.001), *(p<0.008; Holm-Bonferroni-corrected for 6 tests), indicate significant differences between age groups (Wilcoxon ranked-sum tests).

Both midlife and older adults showed a comparable age-related decline in *reward sensitivity* (both Z ≤ -4.02, p < 0.0001; effect size d ≥ 0.6) and *learning rate* (both Z ≤ -2.9, p ≤ 0.003; effect size d ≥ 0.5) compared to young adults, however older adults showed the lowest estimates for *sensitivity to punishment* (Z = -5.69, p < 0.0001; effect size d = 1.0, compared to young adults) as summarized in Figure 3. The modeling approach revealed no significant difference in the *Pavlovian bias across groups* (p = 0.14). For the *irreducible noise* parameter values ranged between 0 and 1 where 1 indicates no distortion to the softmax rule and 0 indicates a flat softmax i.e. random choices regardless of value differences. We observed an age-related difference in the *irreducible noise* parameter (χ^2^ (3) = 28.82; p < 0.0001) whereby values were highest in young adults and lowest in older adults demonstrating younger adults’ performance was more tightly captured by the winning model.

## Discussion

In this study we reveal, that individual performance in a valanced go/no-go task across the lifespan (7-80 years) is influenced by Pavlovian control independent of age. Furthermore, the ability to successfully orthogonalize action and valence was characterized by an inverted U-shape distribution with peak performance observed in young adults. Computational modeling revealed that superior performance in younger adults compared to all other age groups, was attributable to greater sensitivity to outcomes (both to reward and punishment) coupled with a *low action bias.* In contrast reduced performance in children and adolescents was attributable to an increased bias towards action but not increased reward sensitivity. In midlife and older adults, an age-related decline in performance was attributable to a decrease in *learning rate, reward sensitivity* and especially *punishment sensitivity.* Taken together this study reveals novel age-related discrepancies that cannot be probed using typical *go*/reward *no-go*/punishment style paradigms.

This study set out to investigate the influence of Pavlovian control on instrumental learning responses coupled to action and valence using a go/no-go task. Our results revealed that the participants, independently of age, exhibited an influence of Pavlovian control whereby they were better at initiating an action to gain a reward (*go to win*) compared to punishment (*go to avoid losing*) but also withdrawing an action to avoid punishment *(no-go to avoid losing*) compared to gaining a reward *(no-go to win).* The striking asymmetry in performance across conditions observed here is consistent with previous studies using the same task (Guitart-Masip *et al.*, 2012*b*; Cavanagh *et al.*, 2013, Chowdhury *et al.*, 2013*b*; Guitart-Masip *et al.*, 2014; Richter *et al.*, 2014; de Boer *et al.*, 2019). Furthermore, computational modeling in young adults has previously shown this pattern of behavior can be captured by a model incorporating a Pavlovian bias, where the strength of this bias is related to impaired learning of the conflicting conditions: *no-go to win* and *go to avoid losing* (Guitart-Masip *et al.*, 2012*b*). Here we extend this work to demonstrate the same model can effectively capture learning behavior across the human lifespan. Interestingly we observed no significant difference in the *Pavlovian bias* across the lifespan and instead found performance to be influenced by age-related differences in the ability of the instrumental system to learn the appropriate choice (go or no-go) for each fractal image as quantified by differences in *learning rate* and *sensitivity* to both *reward* and *punishment.*

While children and adolescents demonstrated Pavlovian responding consistent with all other age groups, we observed an underlying preference for action responses regardless of valence which goes against the interpretation that there is an overall increase in *reward sensitivity* during adolescence and rather demonstrates this is only true for rewards coupled to action. In fact, our computational modelling show that children and adolescents show decreased *reward sensitivity* when compared to younger adults. As this is the first study to our knowledge to dissociate or orthogonalize action and valence learning in children and adolescents, our results suggest that increased reward sensitivity reported in previous studies (Galvan *et al.*, 2006; Van Leijenhorst *et al.*, 2010; Somerville *et al.*, 2011; Cohen *et al.*, 2016; Palminteri *et al.*, 2016) may be confounded by preference towards action responses. Furthermore these findings highlight the importance of controlling for action tendencies when investigating reward learning in children and adolescents. Previous work in young adults using the same *go/no-go* task has shown that activity in inferior frontal gyrus (IFG), a region known to be involved in action inhibition (Aron *et al.*, 2014), is associated with *no-go* learning and successful instrumental control (Guitart-Masip *et al.*, 2012*b*). Thus protracted development of prefrontal projections to subcortical regions as previously described (Ziegler *et al.*, 2017) may lead to less top-down regulation and increased action bias in children and adolescents.

Reports on changes in instrumental learning across the lifespan have so far focused nearly exclusively on the comparison of young and older adults. Thus, the investigation of midlife adults signifying the transition phase between young and late adulthood has been comparatively neglected. In our study, midlife adults showed significantly compromised performance in *go* conditions compared to younger adults but not to the extent of older adults demonstrating middle age is a pivotal period across the lifespan. The significant age-related decrease in *reward* but *also punishment sensitivity* in midlife adults may reflect the developmental tasks required at this stage of life. In young adults, self-focused goals such as obtaining autonomy or starting a professional career require increased approach motivation and reward responsiveness. However in midlife the focus shifts to taking care or maximizing the benefit of others (e.g. partner, children, ageing parents) and disengaging from unattainable goals allowing for lower approach motivation (Windsor *et al.*, 2012).

In older adults, we found poorer overall performance in all task conditions, but also considerable inability to learn across trials compared to young adults. A study by Schott *et al.*, (2007) suggested that, compared to young adults, older adults exhibited profoundly reduced mesolimbic activation during reward anticipation, although did activate the ventral striatum during reward feedback. Similarly, Chowdhury *et al.*, (2013*a*) demonstrated that older adults do not show a representation of expected value in the ventral striatum when performing a probabilistic reward learning under basal conditions and that an expected value representation was only observed after boosting the dopaminergic system with L-DOPA. Furthermore, older adults performing a probabilistic reward learning task show an attenuation of value anticipation in the ventromedial prefrontal cortex (vmPFC) that predicts performance in the probabilistic learning task (de Boer *et al.*, 2017). These findings suggest that whilst general reward processing may be intact in older adults, they are impaired in learning the predictive value of probabilistic reward cues. In fact, previous studies have shown that reduced learning in older adults is associated with deficits in the integration and updating of reward information when rewards are uncertain and delivered from probabilistic outcomes (Eppinger *et al.*, 2008; Hämmerer *et al.*, 2011). Additionally, Samanez-Larkin *et al.*, (2007) showed a particularly strong age-related impairment of ventral striatal loss anticipation in older adults. Taken together, these findings are compatible with our current observation of decreased instrumental learning in older adults, which was attributable to an age-related decrease in *reward* but *particularly punishment sensitivity* in the learning model.

A striking finding in our data is that the impairment of the instrumental learning in older adults was especially manifest as decreased performance in the go conditions. The dopamine system is involved in generating active motivated behaviour (Niv *et al.*, 2007; Salamone and Correa, 2012) and instrumental learning through reward-prediction errors (Schultz, 2010). Dopamine depletion leads to decreased motor activity and/or reduced motivation to gain rewards (Palmiter, 2008; Salamone and Correa, 2012). Similarly, previous findings using a variant of the same *go*/*no-go* task have shown that L-DOPA administration invigorated instrumental responding regardless of valence (Guitart-Masip *et al.*, 2012*a*). Therefore, an age-related decline in dopaminergic function as previously described (Bäckman *et al.*, 2006; Karrer *et al.*, 2017), could modulate motivation or vigor of actions independently of valence and may explain the overall decrease in *go* performance observed in older compared to younger adults.

Determining the impact of an aging dopaminergic system on performance in the valanced go/no-go task, however, is not straightforward. Most previous studies support the notion that dopamine facilitates the action by valence interaction during learning (see however Guitart-Masip 2014). A study investigating a genetic variant linked to dopamine D2 receptor expression also highlights a modulatory role for genetic variability within the dopaminergic system in individual learning differences of action-valence interactions (Richter *et al.*, 2014). Another study has shown that boosting dopamine with methylphenidate increases the action by valence interaction in participants with high working memory capacity, a proxy for higher dopamine synthesis capacity (Swart *et al.*, 2017). Finally, a recent PET imaging study has shown that the strength of the action by valence interaction scales with the availability of dopamine D1 receptors in the dorsal striatum independent of age (de Boer *et al.*, 2019). Based on this evidence, one would have predicted a decreased Pavlovian bias in older adults. However, learning the correct contingencies of the orthogonalized go/no-go task may critically rely on high-level cognitive functions, thus lifespan differences reported here may also relate to interindividual differences in working memory and long-term memory. These cognitive functions may also be compromised as a result of age-related decline in grey and white matter integrity (Draganski *et al.*, 2011; Samanez-Larkin *et al.*, 2012, Chowdhury *et al.*, 2013*b*; Callaghan *et al.*, 2014; Acosta-Cabronero *et al.*, 2016; Steiger *et al.*, 2016; van de Vijver *et al.*, 2016) which may influence the ability of the instrumental system to learn the task contingencies as indexed by *learning rate, reward* and *punishment sensitivity.* Therefore, the effect of decreased dopamine function on the strength of the Pavlovian system may be shadowed by the effects of an age-related decrease in executive functions or instrumental abilities related to structural decline.

No significant age-related differences in the *Pavlovian bias* was observed across age groups, although a marked increase in the *Pavlovian bias* was observed in older adults. The substantial heterogeneity observed across all age groups also indicates that the strength of the *Pavlovian bias* across the lifespan cannot be entirely accounted for by age. It has been recently argued that an instrumental bias may also influence learning responses coupled to action and valence (Swart *et al.*, 2017) implying a facilitated learning of go responses leading to reward, as well as an impaired unlearning of no-go responses leading to punishment. Thus, it will be interesting in future work to determine how age-related differences in instrumental biases also influence action-valence learning across the human lifespan.

Finally, a general limitation of the study is that we can only infer developmental and age-related differences from a cross-sectional study. These effects should not be assumed to represent underlying causal relationships, nor can we comment on lifespan trajectories. Future longitudinal studies will be needed to address these questions.

Our results demonstrate a dichotomy between prepotent biases that influence learning at either end of the lifespan with a predominant preference for action responses in children/adolescents compared to reduced instrumental learning from both reward and punishment in older age. Collectively, our results emphasize the importance of orthogonally manipulating action requirements and outcome valence to further understand instrumental learning capabilities across different stages of the human lifespan. Such characteristics may underline important evolutionary conserved mechanisms i.e. heightened action learning in adolescents necessary to facilitate active exploration and independence into adulthood or alternatively adaptation to maintain decision-making abilities despite declining learning ability in old age.

## Materials and Methods

### Participants

Overall, 247 individuals between the age of 7 to 80 participated in the current study and were assigned to one of four age groups: *children and adolescents* (age: 7 to 16, n = 69, 48 (69.6%) males), *young adults* (age: 18 to 30, n = 77, 45 (58.4%) males), *midlife adults* (age: 31 to 60, n = 58, 24 (41.4%) males), and *older adults* (age 61 to 80, n = 44, 20 (45.5%) males) using age ranges in line with previously reported lifespan studies (Tymula *et al.*, 2013; Yang *et al.*, 2016). It was ensured either by a standardized telephone interview or personal clinical interview that none of the participants were affected by a present or past neurological or psychiatric illness, alcohol, or drug abuse or were using centrally acting medication. Cognitive abilities were explicitly assessed in children, adolescents and older adults to ascertain they had intact global cognitive performance (for details see SI Materials and Methods). Adult participants were only included if they had finished compulsory education (minimum 12 years). All participants received detailed oral and written information about the study and gave written consent. For minors, informed consent from children and adolescents as well as their parents was required for participation. The study was approved by the local ethics committee of the University of Magdeburg, Faculty of Medicine, and followed the ethical standards of the Declaration of Helsinki.

### Task and Procedure

All participants had to learn which of four fractal cues, preceding an easy visual target detection task, indicated the need (1) to respond to gain a reward (*go to win*), (2) to respond to avoid losing (*go to avoid losing*), (3) to withhold a response to gain a reward (*no-go to win*), and (4) to withhold a response to avoid losing (*no-go to avoid losing*). After display of the fractal cue (1000 ms), participants were presented with the target detection task (1500 ms). During the visual target detection task, participants were presented with a circle either on the right or left side of the screen and had to decide whether they should indicate (*go*) the target side or refrain from pressing a button (*no-go*). For the *go* conditions, they had to emit a button press indicating the side of the target within 1000 ms. Following the circle, participants obtained one of the following feedbacks (1000 ms): a green up-pointing arrow indicating a win of 30 cent in children and adolescents or 50 cent in the adult groups, a red down-pointing arrow indicating a loss of 30/50 cents, or a yellow horizontal bar representing neither win nor loss. Feedback was probabilistic, thus, in the win conditions 80% of correct choices and 20% of incorrect choices were rewarded. In the lose conditions, 80% of correct choices and 20% of incorrect choices successfully avoided loss. Participants were informed and instructed about the probabilistic nature of the task beforehand.

The task consisted of 240 trials (60 trials for each of the four conditions, presented in a randomized fashion in four runs) and lasted approximately 35 minutes. Before the task, participants were asked to complete 10 practice trials in which only the target detection circles were presented to familiarize themselves with the appropriate buttons on the computer keyboard and to obtain an overall feel for the speed of the task without exposure to any of the fractal cues used in the main task. The possible win/loss per trial was 0,50 €. Children and adolescents received reimbursement and reward in the form of gift vouchers (5 euros) for a local shopping center on completion of the task. Adults received the exact amount they won on completion of the task whereas for children and adolescents, earnings were rounded to 5 or 10 euros gain. Stimuli were presented and responses recorded using the Cogent 2000 toolbox (http://www.vislab.ucl.ac.uk/cogent.php) running on MATLAB (Version 2009b; Mathworks).

### Behavioral data analysis

For Behavioral data analysis SPSS Advanced Statistics v21 (IBM Corporation, Armonk, NY, USA) was used. To test whether a Pavlovian bias was evident in all age groups, mean accuracy rates (%) were used in a four-factorial ANCOVA for repeated measures with action (go vs. no-go), valence (win vs. avoid losing) and time (1^st^ vs. 2^nd^ half) as within-subject factors and age group (children and adolescents vs. young adults vs. midlife adults vs. old adults) as between-subject factor, setting gender as covariate of no interest. Independent samples t-tests were performed to compare performance across age groups using the Holm-Bonferroni correction for six tests.

### Reinforcement learning models

We fitted choice behavior to a set of 6 nested reinforcement learning (RL) models incorporating different RL hypothesis. The base model was a Q-learning algorithm (Sutton and Barto, 1998) that used a Rescorla-Wagner update rule to independently track the action value of each choice given each fractal image (*Q*t*(go)* and *Q*t*(nogo)*), with a learning rate (*ε*) as a free parameter. In the model, the probability of choosing one action on trial *t* was a sigmoid function of the difference between the action values scaled by a slope parameter that was parameterized as sensitivity to reward. This basic model was initially augmented with an irreducible action noise parameter also known as a lapse rate (*ξ* (Talmi *et al.*, 2008) and then further expanded by adding a static bias parameter to the value of the go action (*b)*. The model was then augmented by adding a fixed Pavlovian value of 1 to the value of the go action as soon as the first reward was encountered for win cues, and a fixed Pavlovian value of -1 to the value of the go action as soon as the first punishment was encountered for loss cues. This fixed Pavlovian value was weighted by a further free parameter (Pavlovian parameter) into the value of the go action (*π*). Note that this definition of the Pavlovian value is different from the definition in previous studies that have used this task (Guitart-Masip *et al.*, 2012*b*; Cavanagh *et al.*, 2013; de Boer *et al.*, 2019), as model comparison demonstrated it a better fit than a variable Pavlovian value updated on a trial-by-trial basis (see Table 1). The state (action independent) values for each fractal image were updated on every trial using a Rescorla-Wagner update rule with the same learning rate as the update of the action values. Finally, the model including the static action bias and the Pavlovian bias were augmented by including different sensitivities for reward and punishment. Full equations and a description of all considered models are provided in the Supplemental Information.

### Model fitting procedure and comparison

As in previous reports (Huys *et al.*, 2011, Guitart-Masip *et al.*, 2012*b*) we used a hierarchical Type II Bayesian (or random effects) procedure using maximum likelihood to fit simple parameterized distributions for higher-level statistics of the parameters. Since the values of parameters for each subject are ‘hidden’, this employs the Expectation-Maximization (EM) procedure. For each iteration, the posterior distribution over the group for each parameter is used to specify the prior over the individual parameter fits on the next iteration. All six computational models were fit to the data using a single distribution for all participants. This fitting procedure was, therefore, blind to the existence of different groups with putatively different parameter values. Before inference, all parameters except the action bias were suitably transformed to enforce constraints (log for sensitivity to reward and punishment and Pavlovian parameter and inverse sigmoid transforms for learning rate and irreducible noise. Six modeling parameters were extracted for each individual, namely *sensitivity to reward, sensitivity to punishment, Pavlovian bias, action bias, learning rate* and *irreducible noise.* Models were compared using the integrated Bayesian Information Criterion (iBIC) as previously described (Huys *et al.*, 2011, Guitart-Masip *et al.*, 2012*b*). Small iBIC values indicate a model that fits the data better after penalizing for the number of data points associated with each parameter. Comparing iBIC values is akin to a likelihood ratio test (Kass and Raftery, 1995). Note that the iBIC penalizes those versions of the model fit that use four distributions for each parameter. Finally, for each modeling parameter, Kruskal-Wallis tests were conducted to identify age-related differences between groups. Significant interactions were followed up by subsequent group comparisons *using* Wilcoxon rank-sum tests. Holm Bonferroni-correction (p< 0.05 for 6 tests) was used to correct for the effect of multiple comparisons. All tests were performed two-tailed.

## Acknowledgements

The study was supported by the German Research Foundation SFB 779 (TP A03, TP A07, TP A08) and the European Union’s Horizon 2020 Research and Innovation Programme under Grant Agreement No. 720270 (HBP SGA1). Research in the authors’ labs was also supported by the EU/EFRE-funded “Autonomy In Old Age” Initiative of the State of Saxony-Anhalt. MG-M and LdB were supported by a research grant from the Swedish Research Council (VT521-2013-2589) awarded to MG-M. The authors thank Lena Pietro, Carolin Breitling, Anne Hochkeppler, Iris Mann, Catherine Liebeau, Timo Lemme and Maria Watermann for their help during data acquisition. Moreover, we thank Fabian Senner and Marius Keute for their help during preparation of the manuscript.

## Supporting Information

**SI Table 1:** Go/no-go performance across age groups

|  | 7 - 16 vs<br>18 - 30 | 7 - 16 vs<br>31 - 60 | 7 - 16 vs<br>61 - 80 | 18 - 30 vs<br>31 - 60 | 18 - 30 vs<br>61 - 80 | 31 - 60 vs<br>61 - 80 |
| --- | --- | --- | --- | --- | --- | --- |
| **main effect of age group $F_{3,243} = 19.85, p < 0.0001$** |  |  |  |  |  |  |
| overall accuracy | $t_{143} = -5.96,$<br>$p < 0.0001$ | n.s. | n.s. | $t_{133} = 4.13,$<br>$p < 0.0001$ | $t_{119} = 7.36$<br>$p < 0.0001$ | $t_{100} = 2.86$<br>$p = 0.008$ |
| **age group x action $F_{3,243} = 13.07; p < 0.0001$** |  |  |  |  |  |  |
| go accuracy | n.s. | n.s. | $t_{110} = 4.54,$<br>$p < 0.0001$ | $t_{133} = 3.60,$<br>$p < 0.001$ | $t_{119} = 6.11,$<br>$p < 0.0001$ | n.s. |
| nogo accuracy | $t_{143} = -$<br>$6.66,$<br>$p < 0.0001$ | n.s. | n.s. | $t_{133} = 4.02,$<br>$p < 0.001$ | $t_{119} = 6.64,$<br>$p < 0.0001$ | $t_{100} = 2.75,$<br>$p = 0.007$ |
| **age group x valence $F_{3,243} = 2.96, p = 0.033$** |  |  |  |  |  |  |
| Reward | $t_{143} = -$<br>$5.46,$<br>$p < 0.0001$ | n.s. | n.s. | $t_{133} = 3.41,$<br>$p = 0.001$ | $t_{119} = 5.04,$<br>$p < 0.0001$ | n.s. |
| Punishment | $t_{143} = -$<br>$4.78,$<br>$p < 0.0001$ | n.s. | $t_{110} = 3.30,$<br>$p = 0.001$ | $t_{133} = 4.36,$<br>$p < 0.0001$ | $t_{119} = 7.97,$<br>$p < 0.0001$ | $t_{100} = 3.81,$<br>$p < 0.001$ |
| **age group x time $F_{3,243} = 6.01, p = 0.001$** |  |  |  |  |  |  |
| overall learning<br>gain | n.s. | n.s. | n.s. | n.s. | $t_{119} = 4.69,$<br>$p < 0.0001$ | n.s. |
| **age group x action x time $F_{3,243} = 2.69; p = 0.047$** |  |  |  |  |  |  |
| go learning gain | $t_{143} = -$<br>$3.29,$<br>$p = 0.001$ | n.s. | n.s. | n.s. | $t_{119} = 3.07,$<br>$p = 0.003$ | n.s. |
| nogo learning<br>gain | n.s. | n.s. | $t_{110} = 3.0,$<br>$p = 0.003$ | n.s. | $t_{119} = 3.59,$<br>$p < 0.001$ | $t_{100} = 2.94,$<br>$p = 0.004$ |
| go 1<sup>st</sup> half | n.s. | n.s. | $t_{110} = 4.17,$<br>$p < 0.0001$ | n.s. | $t_{119} = 4.24,$<br>$p < 0.0001$ | n.s. |
| go 2<sup>nd</sup> half | $t_{143} = -$<br>$3.57,$<br>$p < 0.001$ | n.s. | $t_{110} = 3.93,$<br>$p < 0.001$ | $t_{133} = 3.92,$<br>$p = 0.0001$ | $t_{119} = 6.69,$<br>$p < 0.0001$ | n.s. |
| nogo 1<sup>st</sup> half | $t_{143} = -$<br>$6.95,$<br>$p < 0.0001$ | $t_{124} = -$<br>$2.90,$<br>$p = 0.004$ | n.s. | $t_{133} = 3.56,$<br>$p = 0.001$ | $t_{119} = 5.33,$<br>$p < 0.0001$ | n.s. |
| nogo 2<sup>nd</sup> half | $t_{143} = -$<br>$5.68,$<br>$p < 0.0001$ | n.s. | n.s. | $t_{133} = 3.33,$<br>$p = 0.001$ | $t_{119} = 6.83,$<br>$p < 0.0001$ | $t_{100} = 3.27,$<br>$p = 0.001$ |

Results from independent samples *t*-tests. Only values from significant results are displayed. Holm-Bonferroni correction was applied for six tests per variable.

**SI Figure 1:**
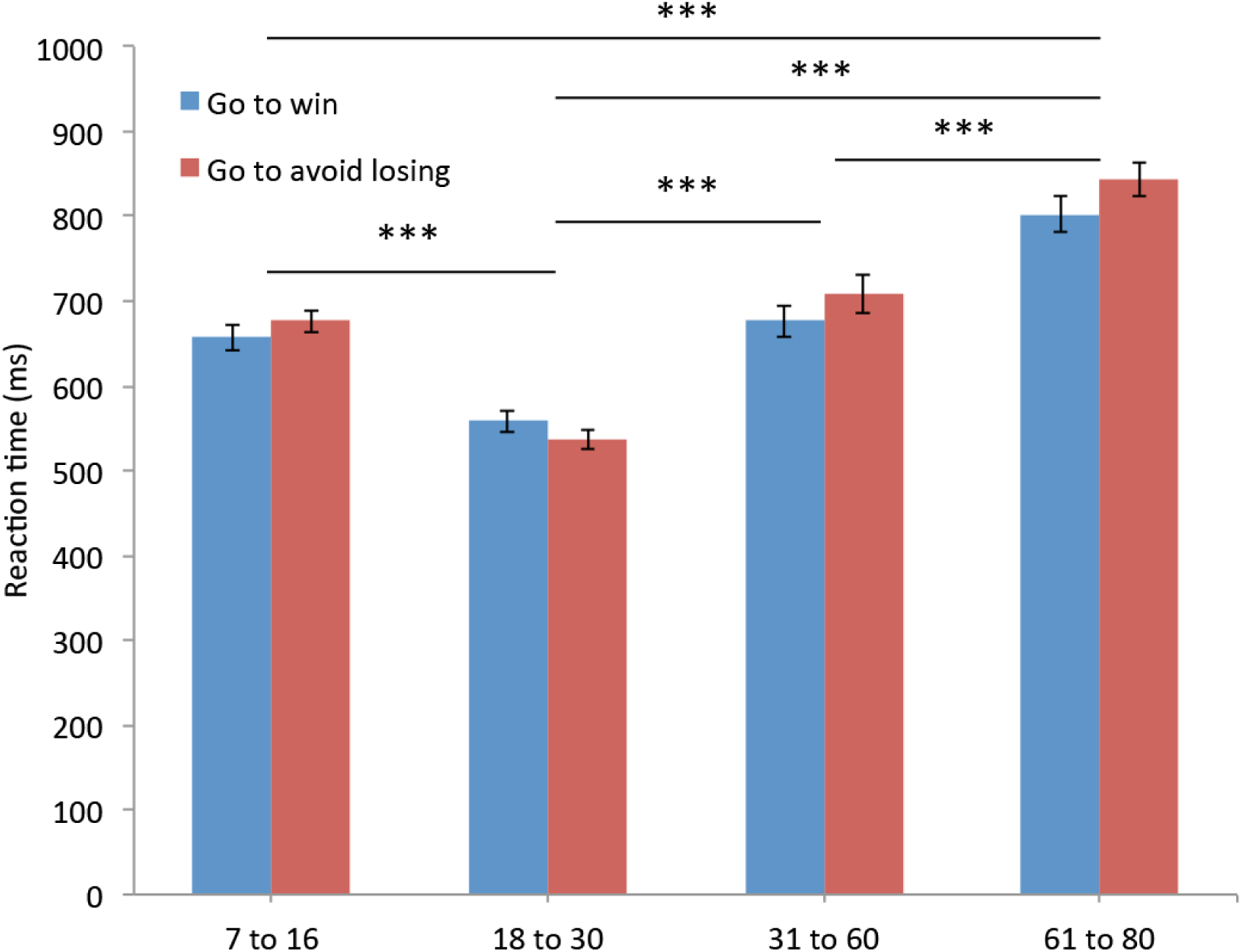
Reaction times for Go responses across the lifespan Mean. (± S.E.M) reaction times (ms) plotted for *go to win* and *go to avoid losing* responses for all age groups. Whilst participants were instructed that the accuracy of their response was more important than speed, a significant difference in *go* response times was observed between groups, whereby young and older adults demonstrated the fastest and slowest responses respectively. ***(p<0.0001) indicates significant differences between groups (1-way ANOVA with Bonferroni *post-hoc* tests).

**SI Figure 2:**
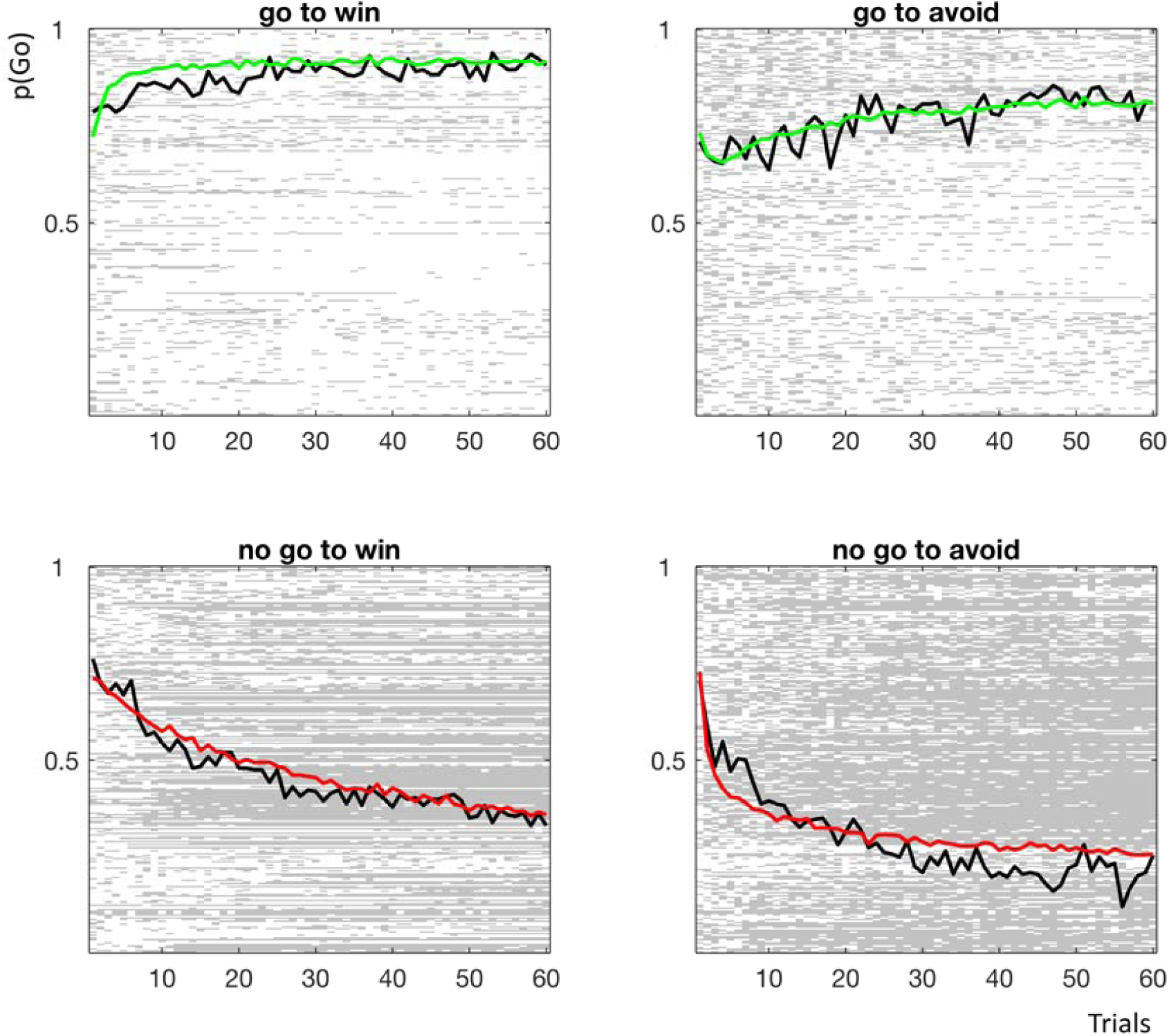
Observed and modeled learning across the lifespan. Modeling parameters from the winning model were used to generate simulated choice data. The simulated group mean probability of performing a go response on each trial is plotted in colored lines (green for go conditions, where go is the correct response; red for no-go conditions, where no-go is the correct response). The mean for all participants’ actual performance is plotted in black lines, reflecting the proportion of actual go responses on each trial. In the plot area, each row represents choice behavior for each participant (n=247) corresponding to a total of 247 pixels per trial. A white pixel illustrates that a participant chose a go response on that trial whilst a gray pixel represents a no-go response.

## SI Materials and Methods

### Assessment of cognitive function and medical history

Participants from the age of 7 to 16 were recruited from a pool of typically developing children and adolescents at the Department of Child and Adolescent Psychiatry and Psychotherapy, University of Magdeburg, or through advertisements in a local newspaper. Participants and their parents were interviewed with the German adaptation (Delmo et al., 2000) of the Revised Schedule for Affective Disorders and Schizophrenia for School-Age Children: Present and Lifetime Version (K-SADS-PL – DSM-IV; Kaufman et al., 1997). Diagnostic tests were selected according to age: Intelligence was assessed using the German adaptation of the *Culture Fair Intelligence Test - Scale 20* (age ≥ 9 years; CFT-20-R; Weiss, 1997) or *Scale 1* (age < 9 years; CFT 1-R, Weiss & Osterland, 2013). *The d2 – Attention Endurance Test* (age ≥ 9 years; Brickenkamp, 2002) or the *bp-Test* (subtest from a developmental test battery for elementary school children; *Basisdiagnostik umschriebener Entwicklungsstörungen im Grundschulalter*; age < 9 years; Esser et al., 2008) were used to measure attentional performance. The Youth-Self-Report (YSR, age > 10 years) and the Child-Behavior-Checklist (CBCL; Achenbach, 1991a,b) were included as further clinical measures. None of the participants received a clinical diagnosis on participation or reported any history of neurological disorders.

All young, midlife and older adults were recruited at the Leibniz Institute for Neurobiology and Institute of Cognitive Neurology and Dementia Research in the University of Magdeburg. All older adults were screened to ensure intact global cognitive performance using a brief neuropsychological battery comprising mini-mental state examination, *Stroop test* in German language and Logical memory test parts I, and II, from the Wechsler memory scale. Individuals with depression were excluded using the Becks Depression Inventory II. Older adults’ alertness and divided attention was assessed using the *Test of Attentional Performance* (TAP). Any individuals known to have had neurological or major psychiatric illness, myocardial infarction, significant cardiovascular history or diabetes mellitus were not eligible for participation.

### Computational modeling of the behavioral data

As in previous experiments where this task has been used (Cavanagh et al., 2013; Guitart-Masip et al., 2012, 2014), we fit six nested models to the observed behavioral data in order to test different instrumental and Pavlovian reinforcement-learning hypothesis. In all models, expected values *Q(a,s*) on each trial t where calculated for each action *a ϵ*{0,1},on each state *s ϵ*{1,2,3,4}, where the actions can be go and no go and states are the four experimental conditions of our task. *Q*(*a,s*) was based on a simple Rescorla-Wagner or delta rule update equation that was implemented each time and outcome was observed:

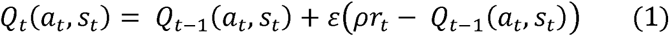

where *ε* is the learning rate. Reinforcements entered the equation through r_*t*_*ϵ*{-1,0,1}and *ρ* is a free parameter that determined the effective size of reinforcements In four models (RW, RW+noise, RW+noise+bias and RW+noise+bias+Pav) there was only one value of *ρ* per subject, meaning that loss of a reward was equally as aversive as obtaining a punishment. The two remaining models (RW(rew/pun)+noise+bias, RW(rew/pun)+noise+bias+Pav) included different values of the parameter *ρ* for reward and punishment trials.

A squashed softmax function (Sutton and Barto, 1998) was used to calculate the probability of selection each action on a given state:

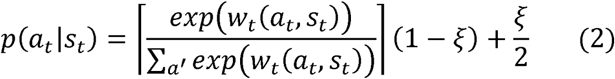

where *w*{*a*_*t*_,*s*_*t*_} reflects the propensity of selecting action *a* in state *s*, and *ξ* is the irreducible noise, which was kept at 0 for one of the models (RW), but was free to vary between 0 and 1 for all other models. Different models varied in the way *w*{*a*_*t*_,*s*_*t*_} was constructed. In the simplest models (RW and RW+noise), *w*{*a*_*t*_,*s*_*t*_}= *Q*_*t*_{*a*_*t*_,*s*_*t*_}

Further models added extra factors to the action propensities. For models that contained a bias parameter, the action weight was modified to include a static bias parameter *b*:

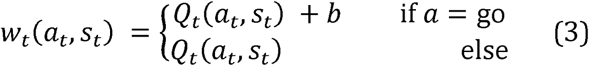

For the model including a Pavlovian factor (RW+noise+bias+Pav and RW(rew/pun)+noise+bias+Pav), the action weight consisted of three components:

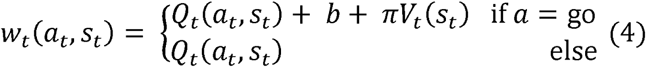

where *π* was again a free parameter. The Pavlovian value V_t_ was determined by the first experienced reinforcing feedback on state *s*. For “win” states, this Pavlovian value was set to 1 after the first trial on which a win outcome was experienced, and for “loss” states, this Pavlovian values was set to -1 after the first trial on which a loss outcome was experienced. Thus, for the ‘avoid loss’ conditions, in which the *V*(*s*) would be non-positive, the Pavlovian parameter inhibited the go tendency in proportion to the negative value *V*(*s*) of the stimulus, while it similarly promoted the tendency to go in conditions in the ‘win’ conditions.

